# Phylogeny, ethnomedicinal use and the distribution of phytoestrogens in the Fabaceae

**DOI:** 10.1101/2025.01.17.633163

**Authors:** Kongkidakorn Thaweepanyaporn, Jamie B. Thompson, Nandini Vasudevan, Julie A. Hawkins

## Abstract

**Aim of the study:** Ethnomedicinal knowledge is a critical resource for drug discovery, and when combined with phylogenetic analysis, it increases the precision of bioprospecting. Phytoestrogens, compounds from plants with estrogenic activity, are commonly found across the Fabaceae family and hold the potential for managing menopause-related symptoms. This study focuses on methods to identify novel sources of phytoestrogens from the Fabaceae.

**Materials and Methods:** We identified 183 Fabaceae species traditionally used as aphrodisiacs or with application to control fertility to create a cross-cultural dataset of ethnomedicinal use. Phylogenetic analysis revealed “hot nodes”—lineages with a higher-than-expected number of species with these traits. The known distribution of estrogenic flavonoids was used to test whether the frequency of phytoestrogen-containing species was higher in “hot nodes”. Additionally, we examined the overlap of aphrodisiac-fertility uses with neurological applications, hypothesising that such species may have bioactive compounds with estrogenic properties. Lastly, the “hot-node” lineages without previously known estrogenic flavonoids were identified.

**Results:** Species in hot nodes were more likely to contain estrogenic flavonoids (21% of species), a major group of phytoestrogens, compared to Fabaceae as a whole (7% of species). Moreover, aphrodisiac-fertility species with neurological applications showed even higher search efficiency, with 62% of such species confirmed to contain estrogenic flavonoids. Furthermore, we identified 43 high-priority hot nodes including several notable genera such as *Delonix* and *Indigofera*.

These lineages might represent promising targets for future studies on phytoestrogens.

**Conclusions:** The results demonstrated the predictive power of combining phylogenetic and ethnomedicinal data to guide the discovery of novel drugs with therapeutic potential for menopause, fertility, and neurological health.

## 1. Introduction

Natural products have been and remain promising candidates for drug discovery (Newman & Cragg, 2020). However, whether natural product research is practicable for drug discovery (Amirkia & Heinrich, 2015), and whether traditional uses in ethnomedicine can guide the discovery of new chemical compounds, remains controversial (Gertsch, 2012; Gurib-Fakim, 2006; Saslis-Lagoudakis et al., 2011; Skirycz et al., 2016; Sucher, 2013). Verpoorte (1998), cited by Verpoorte (2000) and Fabricant & Farnsworth (2001), estimated that 6% of all plant species had been screened for biological activity, and 15% evaluated phytochemically. More recent global estimates are lacking, but one study suggests as many as 58% of plants used in ethnomedicine remain uncharacterised (Souza & Hawkins, 2017). Given the size of the unscreened species pool, devising strategies to target species for evaluation has become an area of research interest (Fabricant & Farnsworth, 2001; Holzmeyer et al., 2020). Of the targeting strategies proposed, ethnobotanically-guided screening has the longest history (Fabricant & Farnsworth, 2001). Strategies incorporating phylogenetic or ecological data, or existing phytochemical and pharmacological data are also becoming established (Pellicer et al., 2018; Saslis-Lagoudakis et al., 2012; Souza et al., 2018). Here we apply phylogenetic methods to ethnobotanical use data, exploring whether they can more efficiently target bioactive plant natural products.

Plant lineages that contain significantly more species with ethnomedicinal use were first referred to as hot nodes for bioprospecting by Saslis-Lagoudakis et al. (2011). Since then, hot nodes have been identified for different groups of medicinal plants from other parts of the world, and at varying taxonomic levels. At the generic level, hot nodes for potential anti-inflammatory compounds have been described for genus *Euphorbia* (Ernst et al., 2016), for species of interest to treat malaria in genus *Artemisia* (Pellicer et al., 2018), and for putative antioxidant and antidiabetic bioactivity for genus *Allium* (Teotia et al., 2024). At a higher taxonomic level, hot nodes in the orchid subtribe Coelogyninae that may show antimicrobial properties were identified based on ethnomedicinal uses (Wati et al., 2021). Geographically-focussed studies have examined cross-cultural patterns between Nepal, South Africa and New Zealand (Saslis-Lagoudakis et al., 2012), whilst others have focussed on the Brazilian Fabaceae (Souza et al., 2018), the Chinese Lamiaceae (Zaman et al., 2022) of whole medicinal floras (South Africa, Yessoufou et al., 2015; Ecuador, Atienza-Barthelemy et al., 2025), or pharmacopoeias (China, Zaman et al, 2021; India, Yao et al., 2023). Global studies include a study of angiosperms to identify hot nodes for psychoactive activity (Halse-Gramkow et al., 2016), for antimalarial properties (Milliken et al., 2021) and cancer treatment (Thompson & Hawkins, 2023). Some of these studies have sought to validate the hot node method, for example, confidence in the hot node method is increased where hot nodes include a higher proportion of plant drugs in clinical trials (Ernst et al., 2016; Pellicer et al., 2018; Souza & Hawkins, 2017), or where there is cross-cultural convergence (Saslis-Lagoudakis et al., 2012). At least one study has used a literature search to show that hot node species have relevant biological activity (Teotia et al., 2024). Pellicer et al. (2018) screened for artemisinin in fifteen species, finding four of seven species from hot nodes and five of eight from outside hot nodes contained artemisinin. Their interpretation was that in this case – where a molecule of interest is common throughout the genus - the hot node approach is not effective. Given the increasing application of the hot node method, further tests of its validity are crucial.

Phytoestrogens (PEs) are plant-derived compounds that have similar functions to estrogen. By binding at the estrogen receptor, estrogen (estradiol, E2) or PEs can activate estrogen-responsive genes, which in turn encode proteins that maintain bone, reproductive health, cognition, and cardiovascular function, (O’Donnell et al., 2007; West et al., 2009).

Consuming one common dietary source of PEs, soybean, can offer a range of health benefits one of which is alleviating the symptoms of menopause (Branca & Lorenzetti, 2005). These symptoms include hot flashes, night sweats, vaginal dryness, mood changes, difficulty sleeping, anxiety and decreased libido (Booyens *et al*., 2022). Several medicinal plant drugs containing PEs are also used to reduce hot flashes and night sweats (Hajirahimkhan et al., 2013), vaginal dryness (Rosa Lima et al., 2014), and cardiovascular disease (Rossouw et al., 2007). The varying interactions of PEs with estrogen receptors suggest that different PEs may have specific functions or roles in various tissues (Ceccarelli et al., 2022; Kiyama, 2022). Because PEs can have both therapeutic and cancer risks (Maggiolini et al., 2002; Umehara et al., 2008; Ye & Shaw, 2019), characterising the diversity of PEs to identify therapeutically optimal molecules is desirable. However, the studies of PEs for postmenopausal symptoms comprise a small number of plants. PEs appear to be distributed throughout the Fabaceae, though most plant sources remain uncharacterised, suggesting there are molecules yet unknown (Dixon, 2004; Rutz et al., 2022). Strategies to identify likely sources of novel PEs are therefore needed.

Here we propose a strategy for identifying potential sources of therapeutically optimal, novel PEs for estrogen-related symptoms. A lack or excess of phytoestrogens, particularly from soybean-based foods, has been shown to suppress sexual behaviour development in both male and female rodents during puberty, suggesting that optimal concentrations of PEs can modulate estrogen-driven behaviours (Khan et al., 2008; Kudwa et al., 2007; Sandhu et al., 2020).

Additionally, chemically isolated PEs such as genistein and daidzein have been shown to produce an anxiolytic-like effect in mice, indicating their potential role in reducing anxiety-related behaviours (Rodríguez-Landa et al., 2009; Zeng et al., 2010). The effects of PEs on socio-sexual behaviour may be mediated through a set of hypothalamic or hypothalamic-linked areas in the brain called the social behaviour network (SBN; Newman, 1999), and applications of plant drugs for neurological symptoms might affect in the same regions (O’Donnell et al., 2007; West et al., 2009). Treatments for menopausal symptoms are very rarely described in ethnobotanical literature, but plants with hormone-modulating properties or those with estrogenic activity may be used as aphrodisiacs or to enhance fertility. Since these applications are directly relevant to sexual behaviour and are often well-documented in traditional medicine, we propose that exploration of aphrodisiac-fertility (AF) as a therapeutic category in ethnomedicine could highlight high-activity PEs that may act predominantly in the CNS. Additionally, neurological applications that regulate CNS activity (Dong & Nao, 2023) may intersect with these therapeutic uses, focusing on plants that have specific effects on the CNS. Species with AF use that also have neurological applications could therefore be of particular interest, as candidates for neuro-selective estrogens.

The Fabaceae is a large and medicinally important plant family. Widely distributed, the family comprises approximately 18, 000 species (Lewis, 2005). Fabaceae plants are rich in alkaloids, flavonoids, saponins, tannins, glycosides, and other phytochemicals that contribute to their medicinal properties (Wink, 2013). Several studies show that the family Fabaceae is over-represented in medicinal floras, indicating that its species are often preferred in traditional medicine (Moerman, 1991; Moutouama & Gaoue, 2024; Saslis-Lagoudakis et al., 2011). The importance of the Fabaceae in traditional medicine is matched by research efforts to characterise species. Many traditional plants in the Fabaceae family have been studied for their pharmacological effects. For example, 71% of Fabaceae genera with traditional uses in Brazil have at least one species that has been characterised (Souza et al., 2018). The species diversity, widespread distribution, numerous reported uses (Souza & Hawkins, 2017), availability of phylogenetic information (LPWG, 2017), and multiple reports of estrogenic compounds within this family (Kiyama, 2017; Kiyama, 2022) have motivated us to focus on this family.

Here we identify species traditionally used for AF purposes and for closely related applications to enhance fertility, and that also have neurological applications. We identify hot nodes using ethnomedicinal data and validate them by comparing them to the known distribution of PEs. The phylogenetic analyses to harness the predictive value of traditional medicine that we present here highlight poorly characterised but ethnomedicinally important lineages that are putative sources of novel PEs.

## 2. Methods

### 2.1 Data collection

Species-level data for flowering plants used as medicine were gathered from recent and comprehensive systematic reviews for Brazil (Souza et al., 2018), China (Zaman et al., 2021), the Greco-Roman Mediterranean (Leonti et al., 2023), the sub-Saharan region of Africa (Ajao et al., 2019) and Thailand (Phumthum et al., 2018).

We compiled a list of aphrodisiac-fertility (AF) species in Fabaceae from these sources by using the search terms ‘aphrodisiac’, ‘sexual intercourse’, ‘libido’, ‘fertility’, and ‘sterilisation’. AF plants are those that stimulate sexual desire. Aphrodisiac use refers to sexual desire within the psychological category. However, aphrodisiacs have also been used in other categories, such as fertility, erectile dysfunction, menstrual disorders, and pregnancy, which fall under the genital system and pregnancy categories. For a more extensive search, we included fertility properties in the search terms because sexual desire and fertility are related to each other (Berger et al., 2016) and estrogen and PEs affected both of sexual desire and fertility (Najaf Najafi & Ghazanfarpour, 2018; Scavello et al., 2019).

Whether the species with AF use had other therapeutic uses was recorded from the original sources and by Google Scholar and PubMed searches. Other uses were classified into ten therapeutic applications (general, blood, digestive, eye, circulatory, muscular, neurological, psychological, respiratory, skin, nutritional, and urinary) according to the ICPC-3 International Classification of Primary Care (van Boven & Ten Napel, 2021).

The list of known PEs (Appendix 1), particularly flavonoids, was obtained by referencing a review on estrogenic flavonoids (Kiyama, 2022). These compounds were then cross-referenced with the LOTUS initiative database (Rutz et al., 2022) using ‘stringdist_left_join’ function from the ‘fuzzyjoin’ package with a maximum difference of two characters between words in R (Robinson et al., 2020) to extract Angiosperm species containing estrogenic flavonoids (Appendix 2).

### 2.2 Phylogenetic analysis

We utilised a large time-calibrated phylogeny of the rosids comprising nearly 20,000 species (Sun et al., 2020), and pruned it to retain only the species in Fabaceae from our data using the ‘keep.tip’ function from the ‘ape’ package in R (Paradis et al., 2004). The final phylogeny included 5,626 (31%) of approximately 18,000 Fabaceae species and 651 (85%).of the 765 Fabaceae genera We used this phylogeny, the list of AF species and the list of species with estrogenic flavonoids in our analyses.

The D statistic was calculated as an estimate of the phylogenetic signal of the AF species. The D statistic was calculated using the ‘phylo.d’ function from the ‘caper’ package in R (Fritz & Purvis, 2010).

The nodes on the phylogeny of Fabaceae that include significantly more species with AF application, referred to here as “hot nodes” (Saslis-Lagoudakis et al., 2012), were identified. We predicted the hot nodes for AF use at the species level using the ‘hot.nodes’ function developed by Molina-Venegas et al. (2020). Hot nodes were considered only if they contained fewer than 100 species, following Halse-Gramkow et al. (2016).

To determine whether screening known AF species or species that belong to AF hot nodes is an efficient bioprospecting strategy, we calculated the percentage of known estrogenic flavonoids that belonged to these groups. We refer to this as “search efficiency” (Souza et al., 2018).

We supposed those AF hot nodes that contained no known estrogenic flavonoids might be the sources of novel estrogenic flavonoids. So, species lists for hot nodes that did not include species that are known as estrogenic flavonoids were created.

The predicted lineages, hot nodes and the phylogenetic distributions of species containing estrogenic flavonoids were visualised using the Interactive Tree of Life v5 (Letunic & Bork, 2021).

## 3. Results

### 3.1 AF species and species with known estrogenic flavonoid

According to the five sources, 183 species belonging to 64 genera were the source of AF medicines. Eight were from Brazil, 122 were from China, seven were from the Graeco-Roman Mediterranean, 28 were sub-Saharan and 19 were from Thailand (Supplementary table 1.) We were able to identify 1317 species that were recorded to produce estrogenic flavonoids, showing approximately 7% of the estimated 18, 000 species of Fabaceae are known to produce estrogenic flavonoids (Appendix 3). Fifty-five (30%) of the species used as AFs were known to have estrogenic flavonoids, and these represented 35 genera; we consider screening AF species to have 30% efficiency (**Figure 1**).

**Figure 1.**
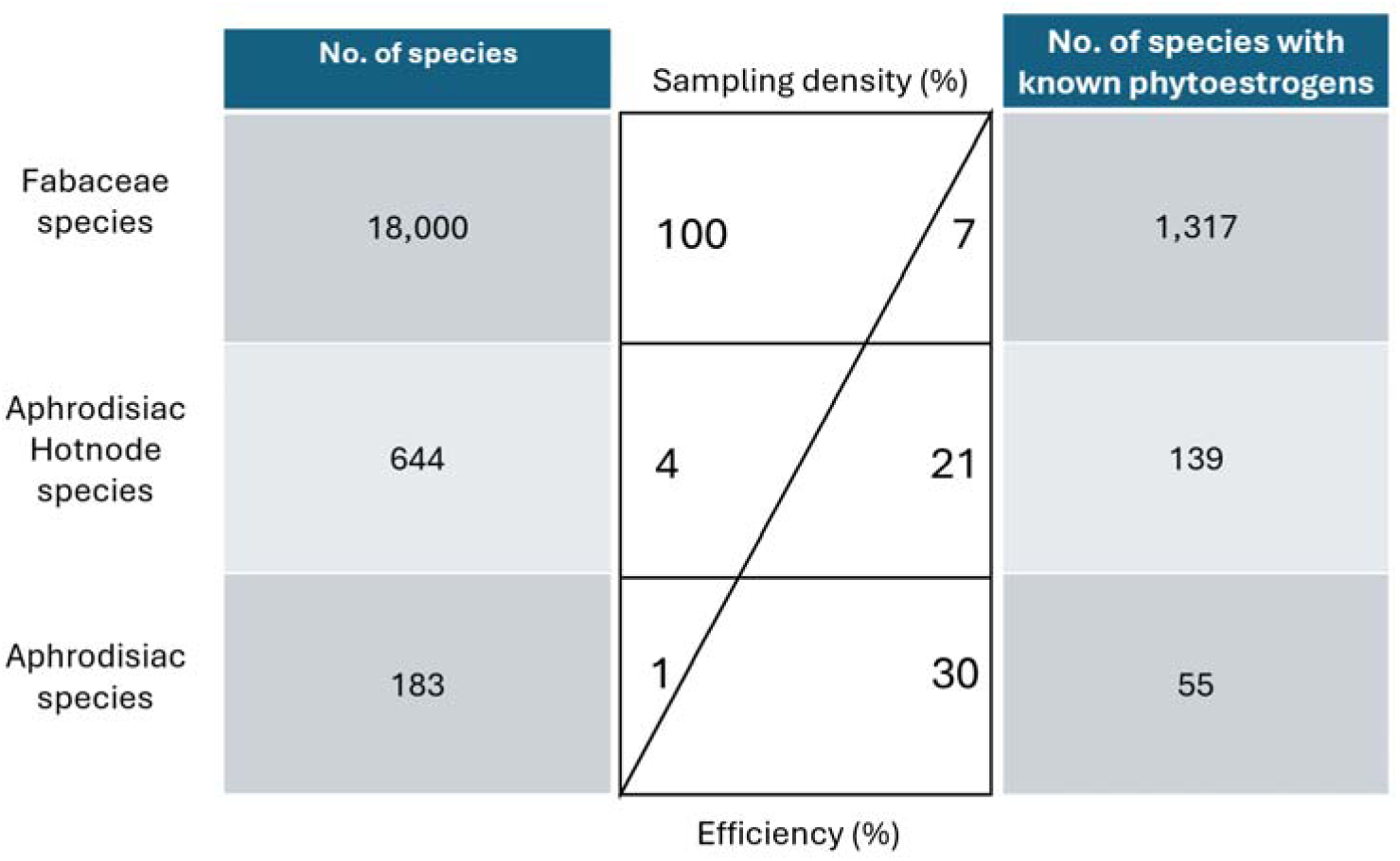
The Comparison of aphrodisiac-fertility species and species that contained estrogenic flavonoid. The first column shows the number of species screened, the central table indicates the sample density and the efficiency of the species with known estrogenic flavonoids by the number of species screened, and the last column shows the number of species with known estrogenic flavonoids.

### 3.2 Predicting lineages with elevated bioprospecting potential

Of the 183 AF species, 106 (57%) were included in the phylogeny. The estimated D statistic for these species was 0.70, indicating a weak to moderate phylogenetic signal for the AF trait. The ‘hot.nodes’ function identified 319 hot nodes. Hot nodes are nested, so our analysis identified 43 highest-level hot nodes (**Figure 2**). These 43 hot nodes comprise 644 species in 142 genera, of which 139 species were known to contain estrogenic flavonoids (21% efficiency; **Figure 1**) The average number of species in the higher-level hot nodes was 29.86 +/-52.27. Of the 43 hot nodes, there were 12 that did not include any species known to have estrogenic flavonoids according to the LOTUS initiative database; the average number of hot node species known to have estrogenic flavonoids was 11.49 species, with a standard deviation of 31.77.

**Figure 2.**
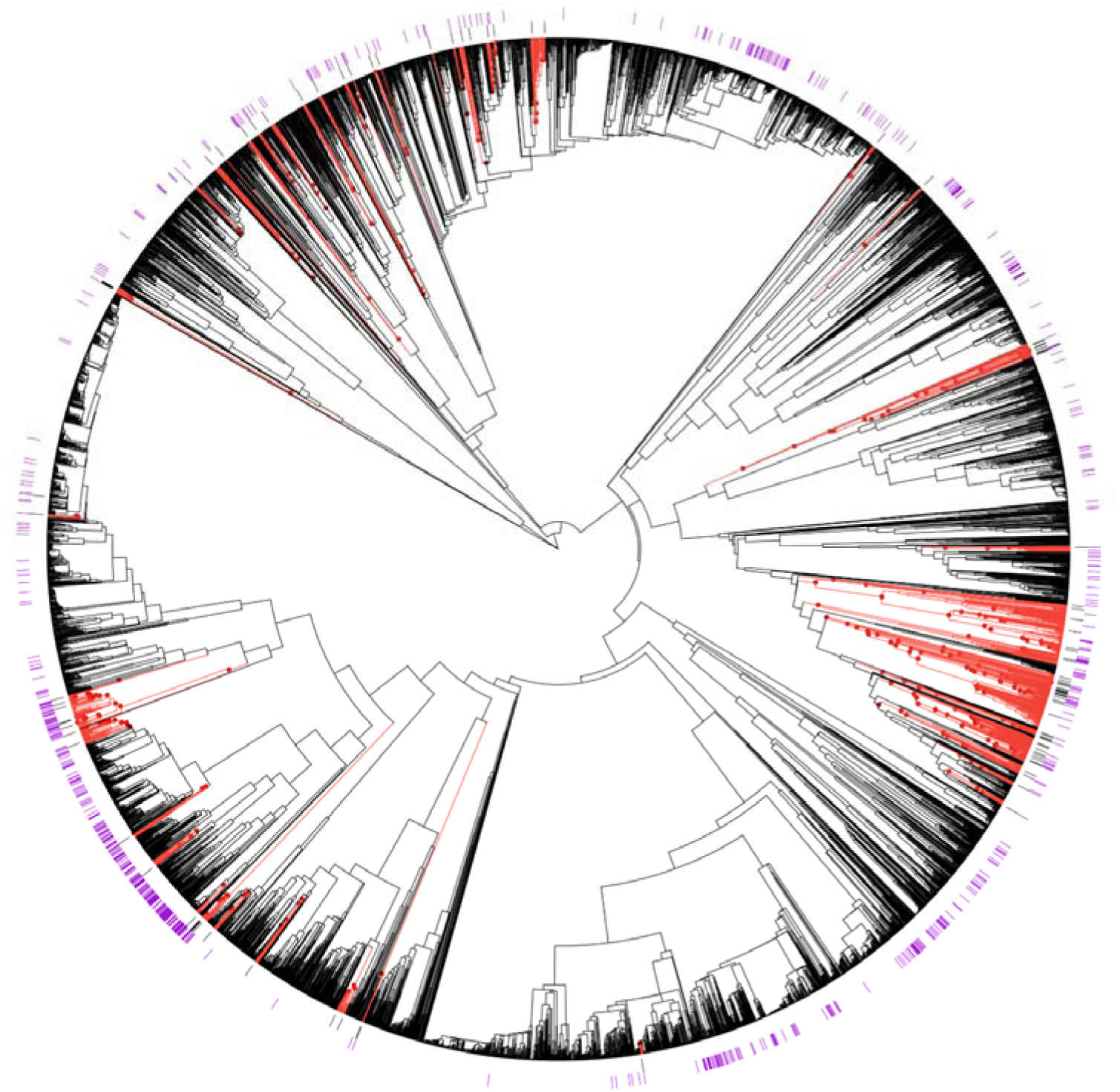
Phylogenetic distributions of species with aphrodisiac-fertility applications and species containing estrogenic flavonoids relative to hot nodes for aphrodisiac-fertility use. species with traditional aphrodisiacs (black bars) and species containing estrogenic flavonoids (purple bars) are indicated on the phylogeny of Fabaceae plants. Hot node lineages for ‘aphrodisiac-fertility’ (red dots) are identified by the ‘hot.nodes’ function developed by Molina-Venegas et al. (Molina-Venegas, Fischer, et al., 2020).

Of the 78 highest-level hot nodes, there were 12 that did not include any species known to contain estrogenic flavonoids. Table 1 shows these hot nodes. The number of species in them ranges from two to 25, with two of the smallest hot nodes only including two species and one node has three species. The first hot node was a sub-family of *Dialioideae* Legume Phylogeny Working Group, and the third cluster contained the genus *Delonix* Raf. The sixth and seventh clusters were in the genera *Vachellia* Wight & Arn., and the eighth cluster included *Senegalia* Raf. and relatives. The ninth cluster was in the genus *Poiretia* Sm. The tenth and eleventh clusters were in the genus *Indigofera* L., while the last cluster was in the genus *Sesbania* Adans.

**Table 1.**
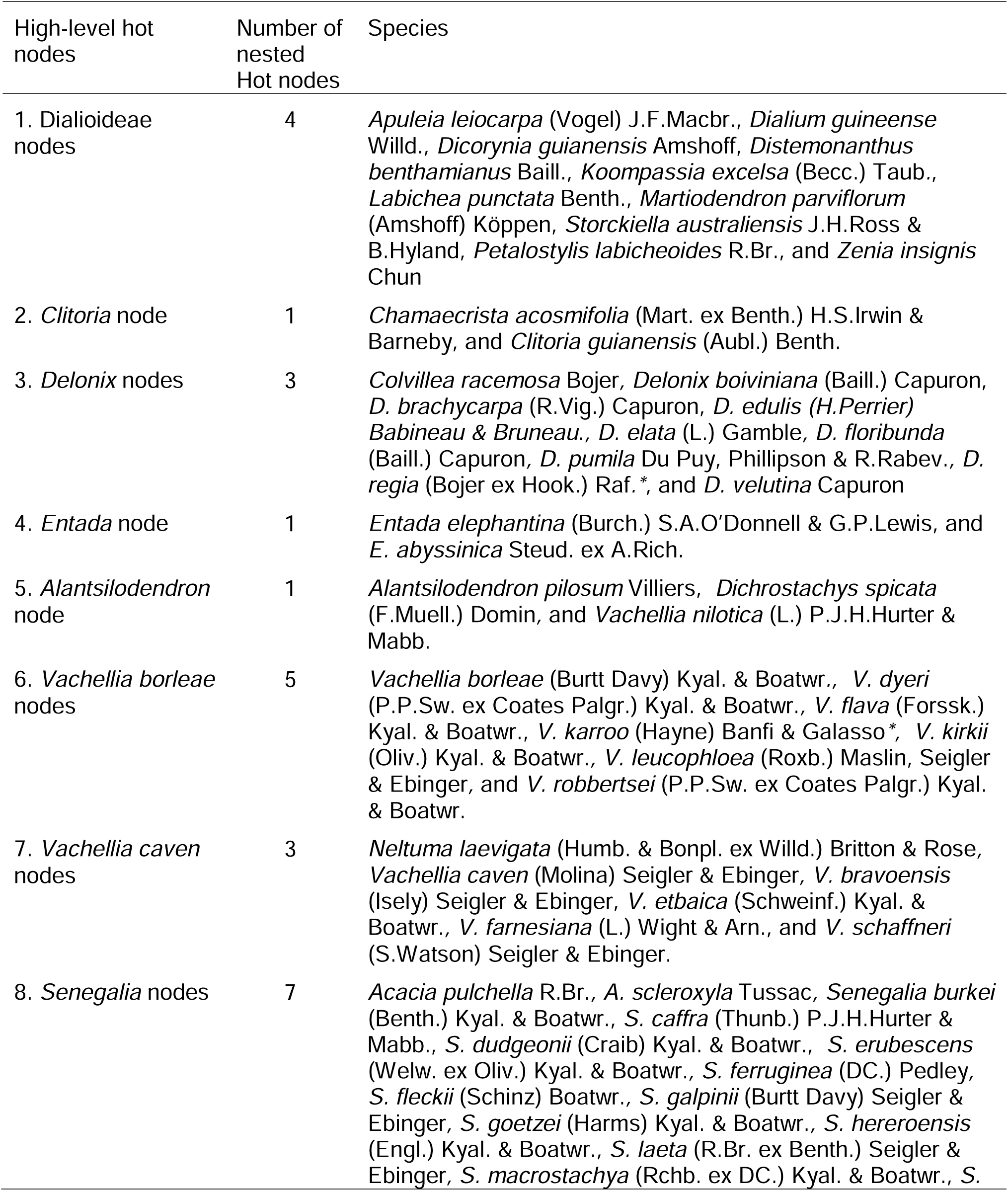
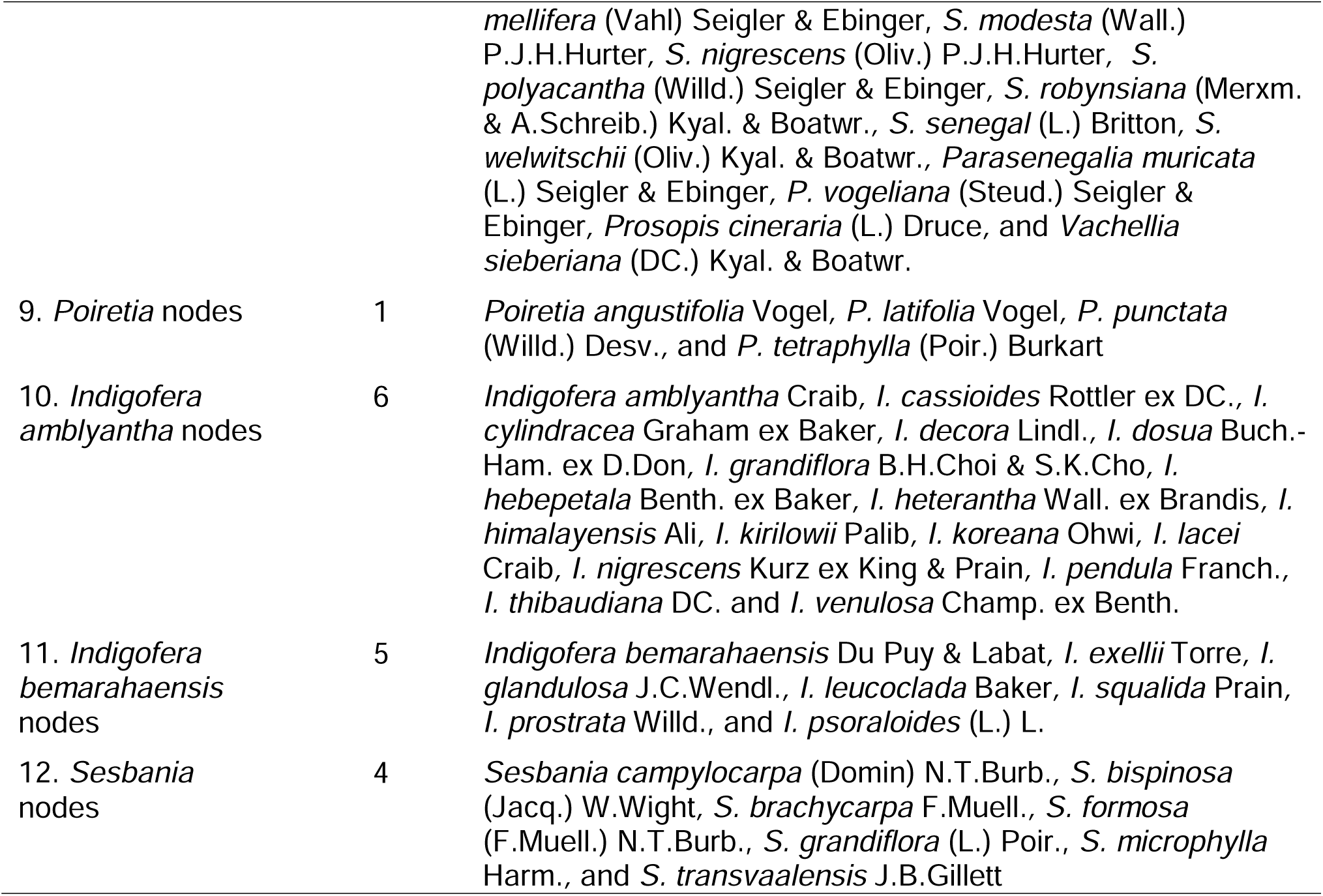
The clusters of hot nodes that include no species recorded as having estrogenic flavonoids in the LOTUS initiative database (Rutz et al., 2022). High-level hot nodes were named by tribe or by most represented genus. If a hot node is repeated in another named node, the descendant node is named alphabetically by the genus appearing first. * Neurological uses.

| High-level hot nodes | Number of nested Hot nodes | Species |
| --- | --- | --- |
| 1. Dialioideae nodes | 4 | *Apuleia leiocarpa* (Vogel) J.F.Macbr., *Dialium guineense* Willd., *Dicorynia guianensis* Amshoff, *Distemonanthus benthamianus* Baill., *Koompassia excelsa* (Becc.) Taub., *Labichea punctata* Benth., *Martiodendron parviflorum* (Amshoff) Köppen, *Storckia australiensis* J.H.Ross & B.Hyland, *Petalostylis labicheoides* R.Br., and *Zenia insignis* Chun |
| 2. *Clitoria* node | 1 | *Chamaecrista acosmifolia* (Mart. ex Benth.) H.S.Irwin & Barneby, and *Clitoria guianensis* (Aubl.) Benth. |
| 3. *Delonix* nodes | 3 | *Colvillea racemosa* Bojer, *Delonix boiviniana* (Baill.) Capuron, *D. brachycarpa* (R.Vig.) Capuron, *D. edulis* (H.Perrier) Babineau & Bruneau., *D. elata* (L.) Gamble, *D. floribunda* (Baill.) Capuron, *D. pumila* Du Puy, Phillipson & R.Rabev., *D. regia* (Bojer ex Hook.) Raf. *, and *D. velutina* Capuron |
| 4. *Entada* node | 1 | *Entada elephantina* (Burch.) S.A.O'Donnell & G.P.Lewis, and *E. abyssinica* Steud. ex A.Rich. |
| 5. *Alantsilodendron* node | 1 | *Alantsilodendron pilosum* Villiers, *Dichrostachys spicata* (F.Muell.) Domin, and *Vachellia nilotica* (L.) P.J.H.Hurter & Mabb. |
| 6. *Vachellia borleae* nodes | 5 | *Vachellia borleae* (Burt Davy) Kyal. & Boatwr., *V. dyeri* (P.P.Sw. ex Coates Palgr.) Kyal. & Boatwr., *V. flava* (Forssk.) Kyal. & Boatwr., *V. karroo* (Hayne) Banfi & Galasso *, *V. kirkii* (Oliv.) Kyal. & Boatwr., *V. leucophloea* (Roxb.) Maslin, Seigler & Ebinger, and *V. robbertsei* (P.P.Sw. ex Coates Palgr.) Kyal. & Boatwr. |
| 7. *Vachellia caven* nodes | 3 | *Neltuma laevigata* (Humb. & Bonpl. ex Willd.) Britton & Rose, *Vachellia caven* (Molina) Seigler & Ebinger, *V. bravoensis* (Isely) Seigler & Ebinger, *V. etbaica* (Schweinf.) Kyal. & Boatwr., *V. farnesiana* (L.) Wight & Arn., and *V. schaffneri* (S.Watson) Seigler & Ebinger. |
| 8. *Senegalia* nodes | 7 | *Acacia pulchella* R.Br., *A. scleroxyla* Tussac, *Senegalia burkei* (Benth.) Kyal. & Boatwr., *S. caffra* (Thunb.) P.J.H.Hurter & Mabb., *S. dudgeonii* (Craib) Kyal. & Boatwr., *S. erubescens* (Welw. ex Oliv.) Kyal. & Boatwr., *S. ferruginea* (DC.) Pedley, *S. fleckii* (Schinz) Boatwr., *S. galpinii* (Burt Davy) Seigler & Ebinger, *S. goetzei* (Harms) Kyal. & Boatwr., *S. hereroensis* (Engl.) Kyal. & Boatwr., *S. laeta* (R.Br. ex Benth.) Seigler & Ebinger, *S. macrostachya* (Rchb. ex DC.) Kyal. & Boatwr., *S.* |
|  |  | *mellifera* (Vahl) Seigler & Ebinger, *S. modesta* (Wall.) P.J.H.Hurter, *S. nigrescens* (Oliv.) P.J.H.Hurter, *S. polyacantha* (Willd.) Seigler & Ebinger, *S. robynsiana* (Merxm. & A.Schreib.) Kyal. & Boatwr., *S. senegal* (L.) Britton, *S. welwitschii* (Oliv.) Kyal. & Boatwr., *Parasenegalia muricata* (L.) Seigler & Ebinger, *P. vogeliana* (Steud.) Seigler & Ebinger, *Prosopis cineraria* (L.) Druce, and *Vachellia sieberiana* (DC.) Kyal. & Boatwr. |
| 9. *Poiretia* nodes | 1 | *Poiretia angustifolia* Vogel, *P. latifolia* Vogel, *P. punctata* (Willd.) Desv., and *P. tetraphylla* (Poir.) Burkart |
| 10. *Indigofera amblyantha* nodes | 6 | *Indigofera amblyantha* Craib, *I. cassioides* Rottler ex DC., *I. cylindracea* Graham ex Baker, *I. decora* Lindl., *I. dosua* Buch.-Ham. ex D.Don, *I. grandiflora* B.H.Choi & S.K.Cho, *I. hebeptala* Benth. ex Baker, *I. heterantha* Wall. ex Brandis, *I. himalayensis* Ali, *I. kirilowii* Palib, *I. koreana* Ohwi, *I. lacei* Craib, *I. nigrescens* Kurz ex King & Prain, *I. pendula* Franch., *I. thibaudiana* DC. and *I. venulosa* Champ. ex Benth. |
| 11. *Indigofera bemarahaensis* nodes | 5 | *Indigofera bemarahaensis* Du Puy & Labat, *I. exellii* Torre, *I. glandulosa* J.C.Wendl., *I. leucoclada* Baker, *I. squalida* Prain, *I. prostrata* Willd., and *I. psoraloides* (L.) L. |
| 12. *Sesbania* nodes | 4 | *Sesbania campylocarpa* (Domin) N.T.Burb., *S. bispinosa* (Jacq.) W.Wight, *S. brachycarpa* F.Muell., *S. formosa* (F.Muell.) N.T.Burb., *S. grandiflora* (L.) Poir., *S. microphylla* Harm., and *S. transvaalensis* J.B.Gillett |

### 3.2 Neurological applications of AF plants

There were 18 of the 165 AF species (10.9%) that also had neurological applications. Of these 18 species, 13 were found in the hot nodes and of those 13 there were eight species (62%) that have been shown to contain estrogenic flavonoids. The eight plants were *Peltophorum africanum* Sond, *Senna siamea* (Lam.) H.S.Irwin & Barneby, *Senna petersiana* (Bolle) Lock, *Mundulea sericea* (Willd.) A.Chev., *Abrus precatorius* L., *Glycyrrhiza glabra* L., *Vicia sativa* L., and *Mimosa pudica* L.. In comparison, only 22% of the 165 AF species without neurological applications found estrogenic flavonoids.

The frequency of other therapeutic applications of the AF medicinal plants is shown in **Figure 3**. 40 AF plants were used to treat ‘general’ disorders, so this was the most common category of use for AF. The second most common category was ‘digestive’ disorders; the ‘neurological’ categories were the next most frequently cited, with 19 AF species reported as used for disorders in each of these categories.

**Figure 3.**
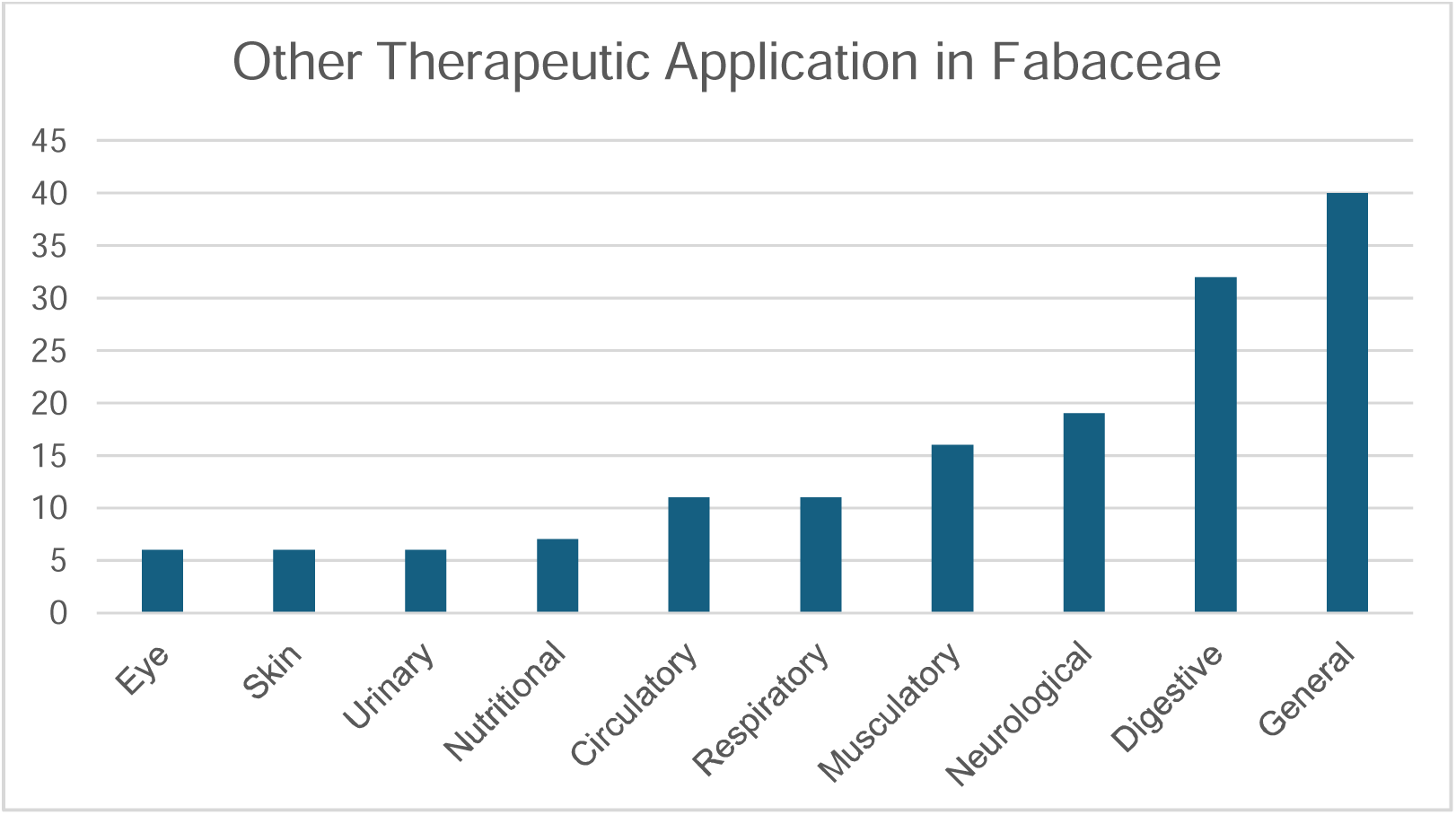
Therapeutic applications of aphrodisiac-fertility plants of Fabaceae. The ten therapeutic categories follow the ICPC-3 International Classification of Primary Care (van Boven & Ten Napel, 2021)

## 4. Discussion

### 4.1 The efficiency of phylogenetic prediction

Therapeutic applications in ethnomedicine have been used in screening programmes to discover new drug leads for many decades (Fabricant & Farnsworth, 2001; Yuan et al., 2016). Species with aphrodisiac and fertility use appear good candidates for the discovery of novel estrogenic flavonoids. A major challenge in drug discovery from plants is the need to select strategically which species to screen, given the impracticality of evaluating all species (Verpoorte, 2000). Effective criteria are essential to identify those most likely to yield useful compounds. Optimisation of the screening process could include focusing on ethnomedicinal species, or plant lineages to which they most frequently belong. This study focused on plants with aphrodisiac and fertility applications in the context of phylogeny to predict lineages with estrogenic flavonoids. Even before we incorporated a phylogenetic framework and hot node analysis, we found that 30% of AF species produce estrogenic flavonoids, compared to 7% of species overall, demonstrating an increased frequency of estrogenic flavonoid-containing species amongst species with AF applications.

Our study reported a D statistic of 0.70, which is a weak to moderate phylogenetic signal for the AF trait (Fritz & Purvis, 2010). However, our community phylogenetic statistics highlighted a pattern of ‘clusters’, since both MPD and MNTP are positive, encouraging us to explore the distribution of estrogenic flavonoids relative to hot nodes for the AF trait. Our finding, that 21% of species in hot nodes have estrogenic flavonoids compared to 7% overall, appears to validate the hot node method, even though the D statistic was suggestive of only moderate predictive power. The search efficiency of 21% for screening hot node species is lower than the search efficiency of 30% for direct screening of AF species. However, hot node species represent more than three times as many candidates for screening without a correspondingly large decrease in known estrogenic flavonoids. We propose that, at least in the case of the Fabaceae, screening hot node species as well as ethnomedicinal species is strategic.

In this study, we show that considering the overlap between a pair of therapeutic applications can enhance the effectiveness of phylogenetic search strategies. The second application we explore here is the application for neurological therapeutic needs. Our data show that when AF applications overlap with neurological uses, these plants are more likely to include estrogenic flavonoids, suggesting a potential dual role in both reproductive and neurological health. We found that 62% of the AF species that are found in hot nodes and that have neurological applications are known to contain estrogenic flavonoids. This is markedly higher than the 22% with phytoestrogens found among the 165 AF species alone. Considering the other therapeutic applications of the AF species, we show use for neurological disorders is the second most common specific application, after digestive applications. This appears to be an elevated frequency, for example in comparison to a ranking of ninth in a study of all therapeutic applications of Brazilian Fabaceae (Souza and Hawkins, 2017), further supporting the view that neurological and AF applications highlight plants with similar bioactivity. Here we highlight the two hot nodes which meet the criteria of hot-node inclusion and neurological use, but which have not been tested for estrogenic flavonoids: *Delonix* nodes and *Vachillia karroo* nodes. A literature survey revealed two species, one from each hot node, *Delonix regia* (Bojer ex Hook.) Raf. and *Vachellia karroo* (Hayne) Banfi & Galasso that did in fact contain estrogenic flavonoids. *Delonix regia* (Bojer ex Hook.) Raf. contains quercetin and its derivatives (Modi et al., 2016) and *Vachellia karroo* (Hayne) Banfi & Galasso epicatechin (Maroyi, 2017); both are estrogenic flavonoids (Kiyama, 2022). Neither species was included in our list of estrogenic flavonoid species because the natural product database was incomplete (Rutz et al., 2022). It would have been possible to make a more complete data set to describe the distribution of estrogenic flavonoids in the Fabaceae for this study. Our study shows that the database has been to sufficient to validate the use of the AF category in hot node analysis. Going forward it is likely that analyses of this kind for other flowering plant families would use the LOTUS initiative database as it is a current and freely available resource.

In our study, we use knowledge of whether plants have estrogenic flavonoids to show that ethnomedicinal uses have predictive power. The presence of estrogenic-flavonoid compounds (Kiyama, 2022) was determined using the LOTUS initiative database (Rutz et al., 2022). Other studies which have sought to validate the hot node method have made comparisons to plant drugs in clinical trials (Ernst et al., 2016; Pellicer et al., 2018; Souza & Hawkins, 2017). Given the increasing application of the hot node method, validation is crucial, so a critical consideration of assumptions related to validation is important. The increase in ‘search efficiency’ we find, from 7% to 30%, is interpreted here as the power of traditional medicine in phytochemical prediction. However, it could also represent a screening bias, because plants used in traditional medicine are more likely to have been investigated and are therefore known to have estrogenic flavonoids. Many of the 18 000 species of Fabaceae have been the focus of phytochemical characterisation, perhaps as many as 43% according to a study of Brazilian Fabaceae that also showed that 52% of the species used in traditional medicine had been the focus of phytochemical or pharmacological study (Souza et al., 2018). However, whilst we recognise this caveat, we do consider the raised search efficiency to validate our method. Three factors in addition to the elevated frequency of species with known PEs are relevant here. Firstly, there is an elevated frequency of a secondary therapeutic application, neurological application, and we had predicted these two uses would be attributed to the same underlying phytochemistry. Secondly, our hot node data are drawn from a cross-cultural sample. This is important because it allows us to discover lineages independently, where cultural beliefs about virility might result in biases in studies of a single culture. Such culture-specific beliefs might be expected for aphrodisiac application, for example, it is well known that bitter tonics are attributed aphrodisiac properties specifically in West Africa and the Caribbean (van Andel et al., 2012). Thirdly, we did find that there were PEs in species we predicted to have them, even when these were not recorded by the LOTUS initiative database. Ultimately, the strongest test of the method may be to assay the plants. Pellicer et al. (2018) screened for artemisinin in fifteen species but did not find that species from hot nodes were significantly more likely to have this bioactive molecule. Whether this is an issue specific to congeneric species, where the biosynthetic pathways needed to produce a bioactive are shared, will be determined by further tests of this kind.

We show that predictive methods of the kind we carry out here merit further investigation. However, the search for therapeutically relevant small molecules has ethical dimensions. The data we analyse here are publicly available data describing ethnomedicinal plant use. Much of these data are available as the result of ethnobotanical research, perhaps motivated by a perceived need to preserve ethnomedicinal knowledge that was experiencing rapid erosion (McManis & Ong, 2018; Schultes, 2007). The ethical dimensions of placing data in the digital commons are now under scrutiny (Mulatinho Simoes & Birchfield, 2024). Where research in this area is carried out by national programmes, in China and India for example, the twin aims of validating and preserving traditional medicine systems can be met, whilst any commercial benefits remain in-country. In our study, the data that we use comes from multiple cultures, and species are highlighted that may not have documented, relevant traditional use. Pellicer et al. (2018) recognise this as an ethical ‘grey area’, as yet not addressed and we further highlight this issue here.

Whether the ethical dimensions of the kind of analysis we present here become the specific focus of rethinking protections for knowledge holders may depend on whether these methods enter the commercial sphere. At present, to the best of our knowledge, work of the kind we present here remains in the academic literature. However, the hot node approach has the advantage of highlighting a broader range of species within the same lineage as known ethnobotanical species. While easily accessible AF plants have been well-characterized in local and regional studies (Ajao et al., 2019; Ganie et al., 2019). Other AF plants remain understudied, perhaps due to the limited availability of plant material. These local and regional studies of local plants could include screening of species that are not used medicinally but that are highlighted by phylogenetic studies. In this way, hot node studies use global data to highlight locally available species to incorporate into local and regional research. Hot nodes provide a better way of identifying species that are a random selection, thus reducing unproductive screening in national programmes.

Even where a wider number of species might be targetted, practical limitations such as the season-dependent chemical composition of plant material, which restrict the time window for recollection, remain. Although many plant-derived natural products have already been isolated and characterized, the amounts available were usually insufficient for extensive testing across a wide range of biological activities (Atanasov et al., 2015). In addition to the accessibility of plant material, the quality was also important. The available plant material often varied in quality and composition, which could hinder the accurate assessment of its therapeutic claims. Chemical composition was influenced not only by species identity and harvest time but also by factors such as soil composition, altitude, climate, processing, and storage conditions (Atanasov et al., 2015). Furthermore, during extraction and isolation processes, compounds could transform and degrade, further complicating the evaluation of their potential therapeutic benefits (Kingston, 2011). These challenges in devising and implementing a screening programme highlight how important it may be to widen the pool of targeted species.

### 4.2 Ethnobotany and phytochemistry of priority hot nodes

In this study, we identified hot nodes that did not include any species known to contain estrogenic flavonoids according to the LOTUS initiative database. These priority hot nodes were investigated in more detail, and several were shown to include estrogenic flavonoids.

The **Dialioideae** nodes include *Apuleia leiocarpa* (Vogel) J.F.Macbr., the bark of this species was used in Peru as a drug to help expel the placenta during childbirth (Odonne et al., 2013), and the root bark of *Distemonanthus benthamianus* Baill. was used for pain relief (Ajibesin et al., 2008). Phytochemical studies of *Apuleia leiocarpa* (Vogel) J.F.Macbr. have revealed the presence of flavones (Braz Filho & Gottlieb, 1971), although their estrogenic activities have not yet been investigated.

The ***Delonix*** nodes includes trees native to Madagascar and East Africa. The most well-known species, *Delonix regia* (Bojer ex Hook.) Raf., has been used in traditional medicine globally and extensively studied for its phytochemical properties (Modi et al., 2016).

Ethnobotanical reports show that its flowers have been used to treat gynaecological disorders (Vidyasagar & Prashantkumar, 2007), and studies have identified flavonoids such as leucocyanidin, cyanidin, and quercetin and their derivatives in the plant (Adjé et al., 2010).

Another species, *D. elata* (L.) Gamble, has been researched for its mosquito-repellent properties *(Govindarajan et al., 2015).* While *D. regia* and *D. elata* (L.) Gamble have been extensively studied, other species within the genus have received less attention.

The genera ***Acacia****, **Senegalia***, and ***Vachellia*** are found in the *Vachellia borleae* nodes*, Vachellia caven* nodes, and *Senegalia* nodes, and were previously grouped as a single genus that was segregated due to its non-monophyly (Kyalangalilwa et al., 2013) These genera are found in Australia, Africa, and other tropical regions (Maslin et al., 2003), and have been widely used in traditional medicine across these areas. Phytochemical investigations have identified flavonoids such as apigenin, catechin, epicatechin, kaempferol, naringenin, quercetin, and myricetin derivatives in species from both Africa and Australia (Subhan et al., 2018). One species, *Vachellia nilotica* (L.) P.J.H.Hurter & Mabb., has been particularly well studied and used to treat a range of conditions, including its use as an aphrodisiac. Research on *V. nilotica* (L.) P.J.H.Hurter & Mabb. indicates that it possesses anti-inflammatory, antioxidant, antidiarrheal, antihypertensive, and antispasmodic properties, in addition to antibacterial, anthelmintic, anticancer, and acetylcholinesterase (AChE) inhibitory activities (Rather et al., 2015).

The ***Poiretia*** nodes consist of twelve endemic species to tropical regions of the Americas. Ethnobotanical reports highlight the use of *Poiretia* species for treating musculoskeletal ailments (Geck et al., 2016). However, some species, such as *Poiretia bahiana* C.Mueller, contain sabinene, a toxic monoterpene (Araújo et al., 2009). On the other hand, *P. bahiana* C.Mueller also contains isoflavonoids with antifungal properties (Araújo et al., 2021). Another species, *Poiretia latifolia* Vogal, contains monoterpenes such as limonene, trans-dihydrocarvone, and carvone, which also exhibit antifungal activities (Nunes Alves Paim et al., 2018).

The ***Indigofera*** nodes are found in the genus *Indigofera*, one of the largest genera within the Fabaceae family (Schrire, 2013) and widely used for medicinal purposes (Gerometta et al., 2020). Several species have been employed as aphrodisiacs, including *Indigofera aspalathoides* Vahl ex DC. in India (Prabhu et al., 2014), *I. cordifolia* B.Heyne ex Roth in Cameroon, Kenya, and Tanzania (Ajao et al., 2019), and *I. flavicans* Baker in Botswana (Ajao et al., 2019). Additionally, *I. cordifolia* B.Heyne ex Roth has been used as an abortifacient in India (Jain et al., 2004) and *I. sanguinea* N.E.Br. in Swaziland (Amusan et al., 2002). Phytochemical studies of various species in the genus have identified numerous flavonoids and isoflavonoids, including apigenin, kaempferol, luteolin, quercetin, genistein, coumestrol, formononetin, and their derivatives (Gerometta et al., 2020).

Finally, the species from ***Sesbania*** nodes are found in tropical and subtropical regions worldwide (Govaerts, 2024). This genus has been used in traditional medicine for treating malaria (Budiarti et al., 2020), dermatology (Mutheeswaran et al., 2011), and headaches (Chellappandian et al., 2012). Phytochemical studies have shown that Sesbania species contain isoflavonoids (Hasan et al., 2012), though much of their potential pharmacological applications remain unexplored.

Despite the documented medicinal uses of many AF species in hot nodes, significant gaps remain in the phytochemical and pharmacological study, particularly regarding their potential estrogenic activities. The *Dialioideae* Legume Phylogeny Working Group subfamily, for instance, has shown promising preliminary results in identifying flavones in *Apuleia leiocarpa* (Vogel) J.F.Macbr., yet its estrogenic potential remains untested. Similarly, while *Delonix regia* (Bojer ex Hook.) Raf. has been extensively studied, other species within the genus *Delonix* Raf. have not received the same attention. This lack of comprehensive research creates a valuable opportunity for further exploration, especially given the known pharmacological relevance of flavonoids. Investigating underexplored genera like *Poiretia* Sm., *Indigofera* L., and *Sesbania* Adams. could yield novel estrogenic flavonoids and other bioactive compounds with potential therapeutic applications.

## CRediT authorship contribution statement

**Kongkidakorn Thaweepanyaporn:** Writing – original draft, review & editing, Methodology, Formal analysis, Data curation, Visualization. **Jamie Thompson**: Writing – review & editing, Methodology. **Nandini Vasudevan:** Writing – review & editing, Methodology, Supervision. **Julie Hawkins:** Writing – original draft, Writing – review & editing, Methodology, Supervision

## Abbreviations

AF: Aphrodisiac-fertility
Pes: Phytoestrogens

## Acknowledgements

We are thankful to Assistant Professor Dr. Methee Phumthum, Mahidol University, for providing Thai ethnobotanical data and advising on this literature.

## Data availability

Data will be made available on request.

## Appendix A. Supplementary data

